# A critical role for CaMKII in behavioral timescale synaptic plasticity in hippocampal CA1 pyramidal neurons

**DOI:** 10.1101/2023.04.18.537377

**Authors:** Kuo Xiao, Yiding Li, Raymond A. Chitwood, Jeffrey C. Magee

## Abstract

Behavioral timescale synaptic plasticity (BTSP) is a type of non-Hebbian synaptic plasticity reported to underlie place field formation in the hippocampal CA1 neurons. Despite this important function, the molecular mechanisms underlying BTSP are poorly understood. The α-Calcium-calmodulin-dependent protein kinase II (αCaMKII) is activated by synaptic transmission-mediated calcium influx and its subsequent phosphorylation is central to synaptic plasticity. Because the activity of αCaMKII is known to outlast the event triggering phosphorylation, we hypothesized it could be involved in the extended timescale of the BTSP process. To examine the role of αCaMKII in BTSP, we performed whole-cell in-vivo and in-vitro recordings in CA1 pyramidal neurons from mice engineered to have a point mutation at the autophosphorylation site (T286A) causing accelerated signaling kinetics. Here we demonstrate a profound deficit in synaptic plasticity, strongly suggesting that αCaMKII signaling is required for BTSP. This study elucidates part of the molecular mechanism of BTSP and provides insight into the function of αCaMKII in place cell formation and ultimately learning and memory.

**Teaser:** The molecular mechanisms of BTSP have been revealed to require the autophosphorylation of CaMKII.

## Introduction

The hippocampus is important for spatial memory in humans and rodents (*1, 2*). During exploratory behavior, hippocampal pyramidal neurons have been observed to fire action potentials at specific locations in the environment(*3, 4*). These “place cells” (PCs) are believed to be formed by behavioral-timescale synaptic plasticity (BTSP), where the weights of active synaptic inputs are bidirectionally altered by a single initiation event—a global voltage signal termed a plateau potential—temporally separated from the to-be modified synaptic input by several seconds (*5*). BTSP has been observed to create new place fields in multiple experimental paradigms including *in-vivo* imaging and whole-cell patch recordings, juxtacellular stimulation, and optogenetic activation of pyramidal neurons in behaving mice (*5*–*12*). BTSP has also been demonstrated in CA1 pyramidal neurons *in-vitro*(*5, 13*). These results suggest that BTSP is a robust synaptic plasticity mechanism for rapidly producing PCs in the hippocampus.

BTSP has been shown to require NMDA receptor and L-type Ca2+ channel activity but other than this the molecular mechanisms involved in BTSP are relatively unknown (*5*). While it seems reasonable to expect that BTSP will share many expression mechanisms with standard LTP, the long time-course of BTSP suggests the necessity of two additional signals. That is, additional molecular mechanisms are needed to temporally filter both the plateau potential (an instructive signal; IS), and synaptic input (an eligibility trace; ET) (Fig. 1A). Although the concept of molecular activity filters (ETs) is nearly 50 years old there is scant evidence concerning the molecules potentially involved (*14*–*16*).

**Figure 1:**
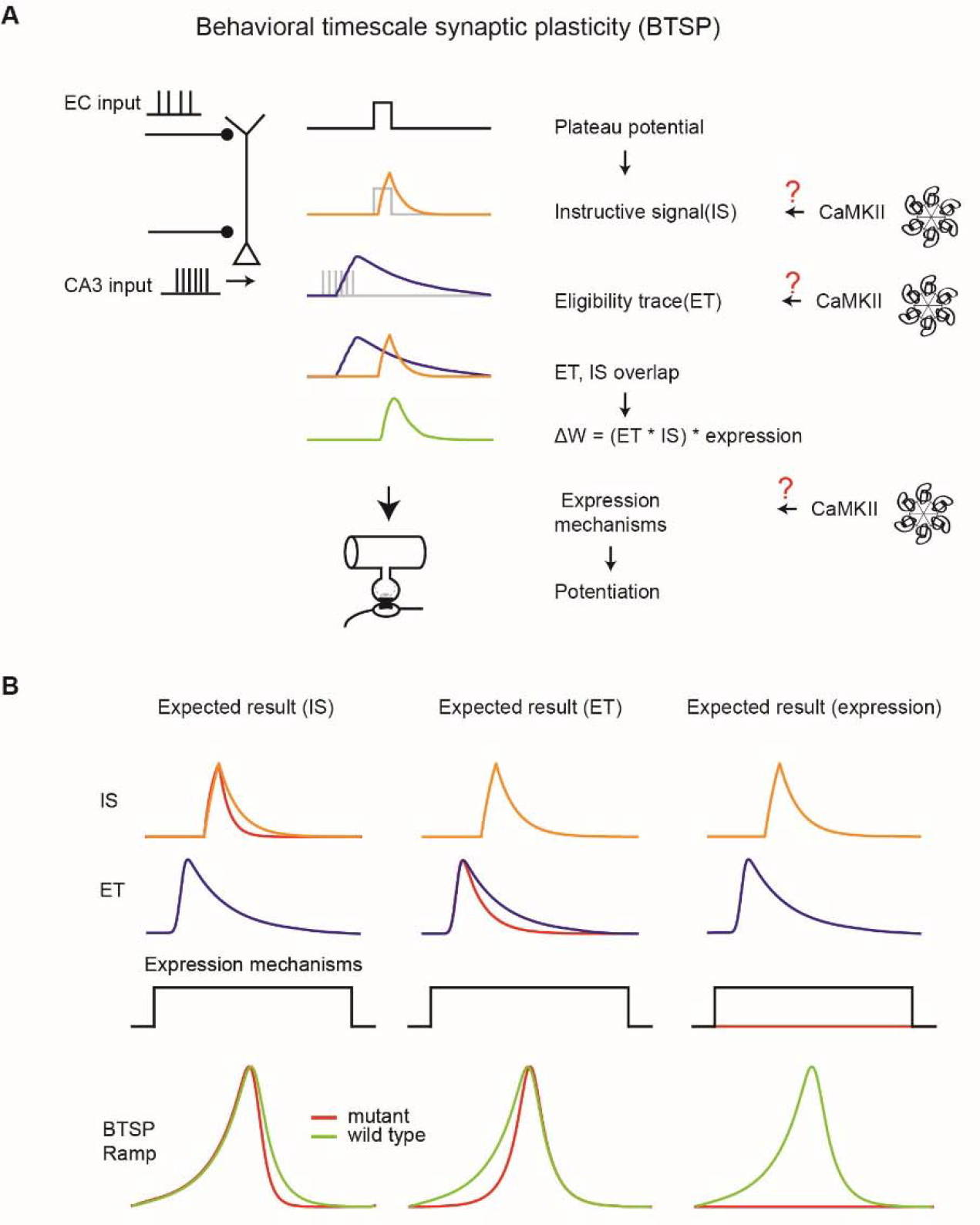
Overview of BTSP and the potential involvement of CaMKII. **(A)**, BTSP model summary. The BTSP model consists of two primary components: an ET and an IS. Eligibility traces are induced by CA3 inputs, while the instructive signal arises from plateau potentials initiated in the tuft region. Gray traces represent electrical events, which produce filtered biochemical signals as shown as colored traces in the schematic. The interaction between eligibility traces and instructive signal determines the delta weight amplitude. We hypothesize that CaMKII plays a role in BTSP, potentially underlying the instructive signal, eligibility trace, or expression mechanism (red question marks). (**B)**, Anticipated experimental outcomes. CaMKII could be implicated in the instructive signal, eligibility traces, or an expression mechanism. Depending on its involvement, distinct effects on the BTSP ramp (depolarization of V_m_ due to BTSP) would be observed: a more asymmetric ramp shape if CaMKII affects the instructive signal, a more symmetric ramp shape if it influences eligibility traces, and the disappearance of the BTSP ramp if CaMKII is involved in the expression mechanism.

Several lines of evidence suggest α-Ca^2+^/calmodulin-dependent protein kinase II (αCaMKII) plays a central role in multiple forms of synaptic plasticity and various properties of αCaMKII identify it as a potential molecular candidate of either the IS or ET (*17*–*20*). The multimeric structure of the αCaMKII holoenzyme endows this molecule with the ability to maintain a phosphorylated state for an extended period (decay time constant 8.2 s (*22*))following transient increases in Ca^2+^ (*21*). At the molecular level, this is mediated by the binding of Ca^2+^/calmodulin (Ca^2+^–CaM) to adjacent regulatory subunits, resulting in the phosphorlylation of T286 in the autoinhibitory portion of the regulatory domain. This disinhibition allows for the kinase activity to persist beyond the initial binding of Ca^2+^–CaM (*22*). Therefore, T286 autophosphorylation allows for the integration of transient Ca^2+^ signals and their transformation into long lasting αCaMKII activation (*23, 24*). The substitution of Thr^286^(T) for Alanine (A, T286A) of αCaMKII has been demonstrated to prevent CaMKII constitutive activity and animals engineered to express this mutation perform poorly in spatial memory tasks, display deficits in hippocampal synaptic plasticity and reductions in spatial selectivity and stability of PCs(*18, 19*). Recent studies using Förster resonance energy transfer (FRET)-based CaMKII sensors revealed that this mutation greatly increases the decay of αCaMKII activation (decay time constant 1.9 s (*21*)) relative to the native autophosphorylated state(*21, 22*)

Based on the above studies, we hypothesize that αCaMKII activity is involved in BTSP. If αCaMKII activity functions as either an ET or an IS, the more rapid decay kinetics resulting from the T286A mutation should decrease the duration of these signals (Fig 1B). This reduced time course would significantly alter the shape of the underlying subthreshold membrane potential (V_m_ ramp) produced by the BTSP potentiated synapses with the exact alteration dependent on which signal (IS or ET) is mediated by αCaMKII. Alternatively, αCaMKII signaling might be a component of the myriad synaptic plasticity expression mechanisms and, in this case, the modified kinetics resulting from the T286A mutation would affect the level of BTSP induction perhaps to the point where BTSP is unable to produce PCs or even any associated subthreshold V_m_ changes (Fig 1B). Finally, if αCaMKII signaling is not involved in BTSP, the T286A mutation would have no effect. Thus, the T286A αCaMKII mutation can aid in teasing apart the specific role of this molecule in BTSP.

To test the effect of faster αCaMKII decay kinetics on BTSP, we performed whole-cell patch-clamp recordings in awake, behaving T286A^+/+^ (Homo) mice, as well as T286A^+/-^ (Het) and wild-type (WT) mice as controls. We found that the BTSP induction in the T286A^+/+^ group resulted in very weak or no BTSP, whereas the Het group and the WT group have similar BTSP. We also observed an increased propensity for CA1 neurons in the T286A^+/+^ group to fire spontaneous plateau potentials, suggesting alterations in excitability either in CA1 or other regions of the hippocampal network. Interestingly, these spontaneous plateaus in the homozygous group also did not produce PC activity. To control for the possibility that the above effect of T286A mutation resulted from an alteration in the hippocampal network such that CA1 received only inconsistent afferent input, we tested BTSP induction in hippocampal brain slice, where we can reliably and consistently activate a given set of synaptic inputs. Consistent with the *in-vivo* results, BTSP induction here produced minimal potentiation in the T286A slices relative to control. In addition, standard in vitro measures of membrane excitability did not reveal differences that would likely contribute to the increase in spontaneous plateau potentials observed during behavior. Together, our results reveal that αCaMKII plays a central role in BTSP at some stage other than in the generation of ETs or ISs, suggesting that αCaMKII signaling is likely required for the synaptic plasticity expression process.

## Results

### Experimental design and behavioral quantification

To determine the effect of altered αCaMKII signaling on PC formation, we performed whole-cell recordings from CA1 pyramidal neurons in awake behaving mice. Animals were well acclimated to the experimenter and trained to run on a treadmill to receive a water reward at a fixed location (Fig 2A). The treadmill was a 180 cm belt demarked with distinct visual and tactile cues on its surface (Fig 2B). As described previously (*25, 26*) well-trained animals exhibited stereotyped patterns of running and licking as they neared the reward location (Fig 2C–G). The fraction of licks within a 15 cm zone centered around the reward location were similar in the WT and Het groups but significantly reduced in the Homo group (Fig 2C, D). The homozygous animals typically displayed faster running speed across the track and the minimum speed at the reward site was faster compared to the WT and Het groups (Fig. 2E–G). Despite these quantitative differences, the characteristic increases in licking and slowing near the reward zone indicated that the homozygous animals engaged in the task and could be directly compared to the WT and Het groups.

**Figure 2:**
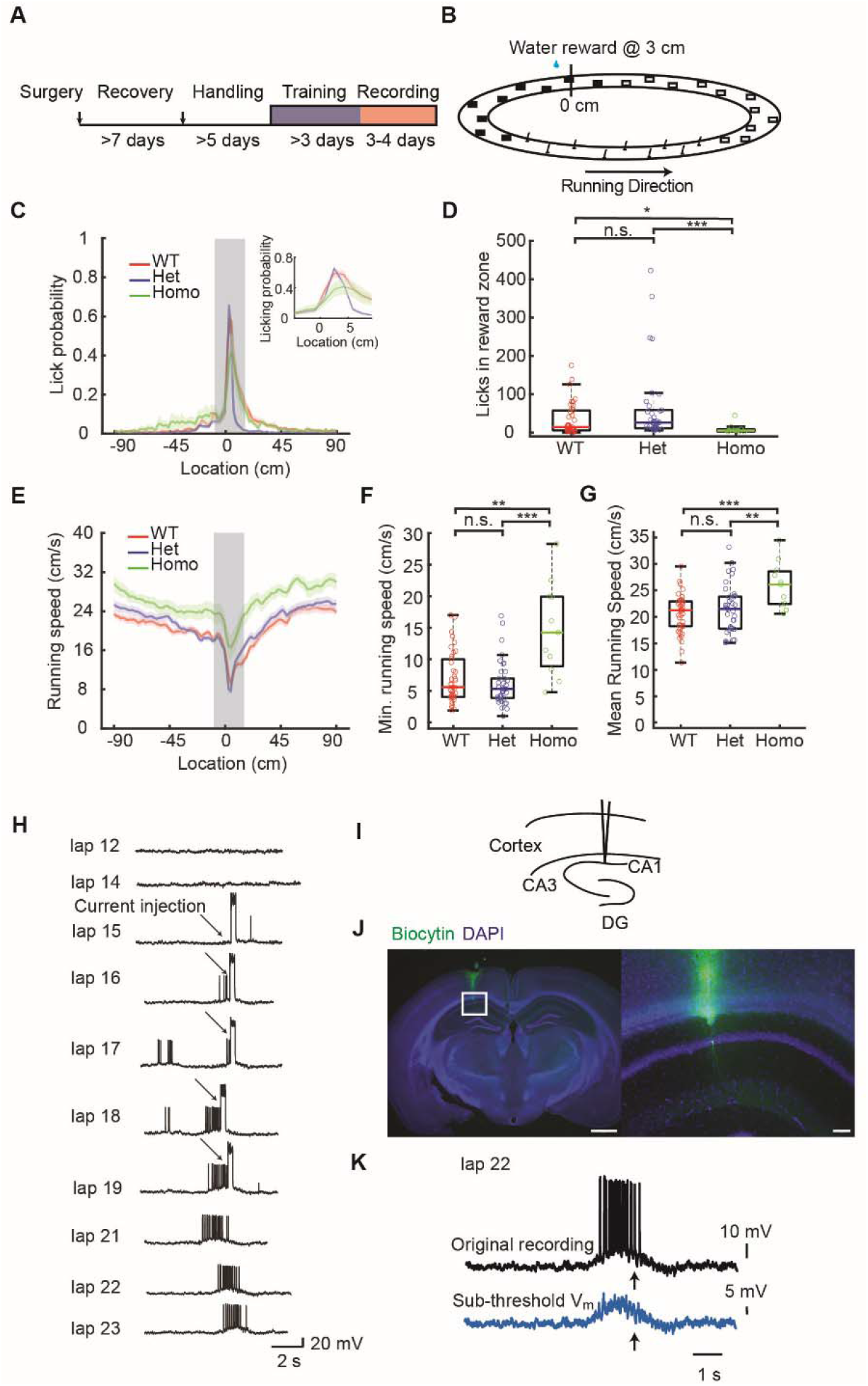
Experimental set up, quantification of animal behavior for BTSP induction and example BTSP recordings. **(A)**, Experimental procedure involving surgery, recovery (7 days), handling (5 days), and training (minimum of 3 days) prior to recording sessions. (**B)**, Linear track configuration with cues distributed in various sections; reward site at 3 cm. (**C)**, Licking probability at different spatial locations for WT, Het, and Homo groups, with reward location at the center (3 cm). Insets display an enlarged view of the region surrounding the reward site. Lines with shaded backgrounds represent mean ± SEM. (**D)**, the relative licking probability increase in the reward zone (see methods). P-values: 0.131 (WT vs. Het), 0.0414 (WT vs. Homo), 4.71e–4 (Het vs. Homo). (**E)**, The running speed in different spatial locations. Data are from all the laps recorded averaged. (see methods, reward location in 3 cm). Lines with shaded backgrounds represent mean ± SEM. (**F)**, Minimum averaged running speed value for all recorded laps per recorded cell. P-values: 0.774 (WT vs. Het), 0.00305 (WT vs. Homo), 3.58e–4 (Het vs. Homo). (**G)**, mean value in averaged running speed of all recorded laps corresponding to each cell. P-values: 0.979 (WT vs. Het), 8.00e–4 (WT vs. Homo), 0.00705 (Het vs. Homo). (**H)**, Example cell exhibiting BTSP induction process, where a place field is induced by BTSP in a previously silent CA1 neuron. Arrows indicate current injections. Scale bars: 20 mV, 2 s. (**I)**, Schematic of in-vivo recording site. (**J)**, Post hoc histology example of recorded CA1 neuron. Right panel provides an enlarged view of the left panel. Blue color: DAPI; Green color: Biocytin. Scale bars: 1000 μm (left), 100 μm (right). (**K)**, example of the process to calculate the subthreshold voltage depolarization in a place field for *in-vivo* recordings. Subthreshold membrane potential (blue) was extracted from original V_m_ recording as described previously (*5*). Parametric statistical analysis utilized one-way ANOVA with Bonferroni correction; non-parametric analysis employed Kruskal-Wallis test with Dunn’s test for multiple comparisons. *P< 0.05, **P< 0.01 and ***P< 0.001; NS, not significant (P≥ 0.05). Sample sizes: n=39 (WT), n=38 (Het), n=11 (Homo).

As described previously, BTSP can be reliably induced by strong somatic depolarization that evokes plateau potentials to establish location-specific firing (*7, 12, 27, 28*). In WT animals, CA1 pyramidal neurons are typically silent, but brief large-amplitude current injections (300 ms, ∼700 pA) at a defined location will transform silent cells into PCs. Although our standard induction protocol utilizes 5 repetitions of somatic current injection, the location-specific firing of APs often develops after the first induction trial and grows more pronounced with subsequent trials (Fig 2H–J). The AP output of this newly formed PC begins at a location preceding location of induction presumably because synaptic inputs active at these locations have been potentiated (*5*). Indeed, when the membrane potential is filtered to include only subthreshold values the underlying V_m_ ramp begins at locations preceding the location of induction (Fig 2K) and is prospectively asymmetric.

### The V_m_ ramp is significantly reduced in the homozygous T286A animals

Because αCaMKII signaling acts to temporally filter the postsynaptic changes in V_m_ and associated Ca^2+^ influx in a manner consistent with the molecular underpinning of either an ET or IS and the T286A mutation is known to hasten the decay of this signaling and adversely affect synaptic plasticity, PC stability and spatial memory, we compared the V_m_ ramp resulting from a standard BTSP induction protocol in all groups of animals. In the WT and Het groups, comparisons of the subthreshold V_m_ before and after BTSP induction revealed the instantaneous formation of depolarizing synaptic responses for many seconds around the induction location (90 cm). However, little or no such changes in subthreshold V_m_ were observed in the Homozygous group (Fig. 3A). A further comparison of the average subthreshold V_m_ for the 5 laps before and after BTSP induction reveals a characteristic asymmetric V_m_ ramp for the WT and Het groups, that peaked at 8.0 ± 0.54 mV and 7.6 ± 0.62 mV respectively. In contrast, the subthreshold V_m_ in the Homozygous group was minimally increased by 2.3 ± 0.75 mV following BTSP induction (Fig. 3B–G). The evolution of the V_m_ ramp during the induction protocol was similar for the WT and Het groups, exceeding 8 mV within 5 laps after the start of BTSP induction while the change in Vm in the Homo group was < 2mV (Fig. 3H). To control for the possibility that somatic depolarization might be ineffective at reproducing the conditions of a naturally occurring plateau potential in the Homozygous group, we also compared the subthreshold V_m_ before and after spontaneous plateau events and found these were also ineffective at producing V_m_ ramps in these mice (Supp. Fig. 1)

**Figure 3:**
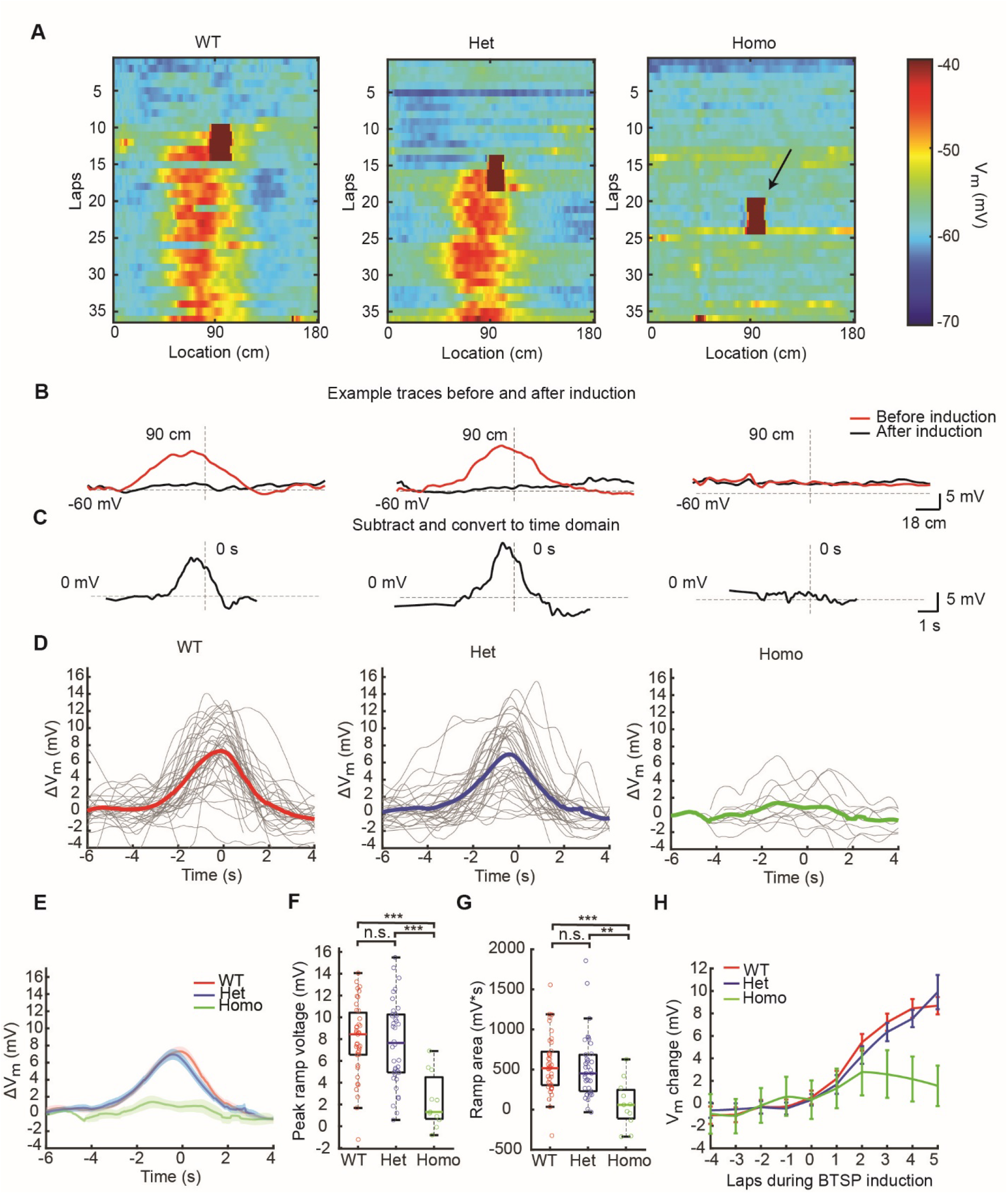
Reduced BTSP *in-vivo* in CaMKII T286A homozygous mutant mice. **(A)**, Color plot of three example cells from WT, Het, and Homo groups, displaying subthreshold membrane potential for each lap in the ∼180 cm track. Arrowheads indicate plateau potentials during BTSP induction events. (**B**–**C)**, Example traces showing the ramp post-BTSP induction for WT, Het, and Homo groups. The subthreshold Vm before and after BTSP induction was averaged (5 laps before and 5 laps after), and the resulting ramp is obtained by subtracting the pre-induction trace from the post-induction trace. The subtraction trace is conservatively converted into time domain based on the fastest running speed during induction laps, as described by (*5*). Dashed lines indicate –60 mV and 0 mV voltage, as well as 90 cm location and 0 s time. (**D)**, BTSP ramps for each cell in WT (left), Het (middle), and Homo (right) groups; thicker lines represent group averages. (**E)**, Averaged data for all three groups plotted together, with lines indicating mean ± SEM. (**F)**, Quantification of ramp peak in the three groups. P-values: 1.00 (WT vs. Het), 2.35e–5 (WT vs. Homo), 7.69e–5 (Het vs. Homo). (**G)**, Quantification of the ramp area for the three groups. P-values: 0.854 (WT vs. Het), 4.07e–4 (WT vs. Homo), 0.00262 (Het vs. Homo). (**H)**, Speed of BTSP ramp formation, depicting subthreshold membrane potential in the place before (5.4 cm before) the plateau induction for different laps (5 laps pre-induction and 5 laps during induction). Statistical analysis employed one-way ANOVA with Bonferroni correction. *P< 0.05, **P< 0.01 and ***P< 0.001; NS, not significant (P≥ 0.05). Sample sizes: n=39 (WT), n=38 (Het), n=11 (Homo).

### Slice experiments control for the reliability of synaptic input

The plasticity observed following BTSP induction is believed to be the result of increases in synaptic weight of spatially tuned inputs reliably activated at specific locations as the animal traverses the environment. Because the effect of the T286A mutation may have produced alterations to the circuit providing these inputs, it is possible that the reliability of these inputs being consistently active at a given location may be diminished to the point where it would be unlikely to observe plasticity. To control for this, we next attempted to induce BTSP in the slice preparation where the population of synaptic inputs activated by extracellular stimulation would be more consistent on a trial-to-trial basis (Fig. 4A). To reliably elicit plateau potentials, we utilized a Cs^+^-based internal solution as described previously(*5*). Consistent with previous reports, BTSP induction at the 0-ms interval produced an immediate 3.31 ± 0.508-fold increase in synaptic weight relative to baseline in slices from WT animals. In contrast, slices from the Homozygous animals exhibited very weak BTSP with a modest 1.07 ± 0.119-fold increase relative to baseline (Fig 4B–F, Supp. Fig. 2). BTSP induction did not produce changes in input resistance or paired-pulse facilitation relative to baseline in either group (Fig. 4 G, H). These results add further support to the in-vivo findings suggesting a critical role for αCaMKII in BTSP, that is not related to the filtering of either the synaptic input (i.e.ETs) or the plateau potential (i.e. ISs).

**Figure 4:**
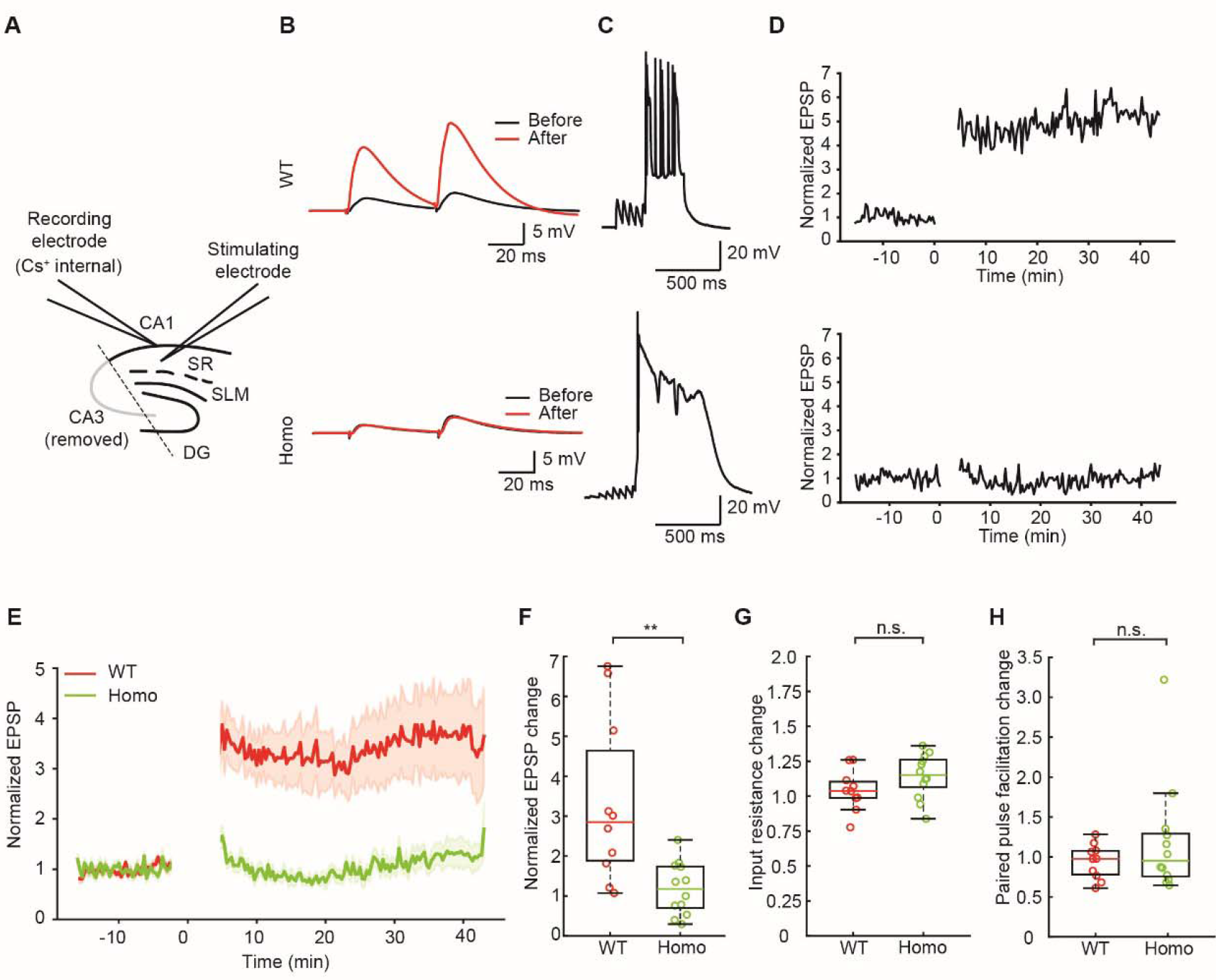
Reduced BTSP in CaMKII T286A homozygous mutant mice in slice experiments. **(A)**, Schematic of the slice experiment, with EPSPs elicited by electrical stimulation in the SR area of the hippocampus. CA3 region was removed to prevent excessive input from CA3 neurons interfering with recordings. (**B)**, example raw traces of EPSPs before and after BTSP induction in WT group (top) and in Homo group (bottom). (**C)**, example traces during the BTSP inductions from WT group(top) and Homo group(bottom). (**D)**, EPSP amplitude plotted against time, with cells recorded for over 15 minutes pre-induction to ensure stable baselines. Post-induction recordings last up to 40 minutes. (**E)**, group data for EPSP amplitude plotted against time, with lines and shaded backgrounds representing mean ± SEM, respectively. (**F)**, quantification of the EPSP amplitude change. the change is calculated by dividing the averaged EPSP amplitude before induction from the EPSP amplitude after induction in each of the cells. P-value: 0.00291. (**G)**, quantification of the changed input resistance (pre- and post-induction) of WT and Homozygous groups. P-value: 0.144. **(H)**, quantification of the changed paired-pulse facilitation rate (pre- and post-induction) of WT and Homozygous groups. The paired-pulse facilitation rate was calculated by dividing the first EPSP’s amplitude from the second EPSP’s amplitude. P-value: 0.295. All statistical analyses employed a two-sample Student’s t-test. *P< 0.05, **P< 0.01 and ***P< 0.001; NS, not significant (P≥ 0.05). n=10 for WT, n=12 for Homo.

### Cellular excitability

Independent of BTSP induction, the spiking behavior of CA1 pyramidal neurons in the T286A animals was different with regards to the probability and duration of spontaneous plateau events (Fig. 5A). Indeed, the duration of these spontaneous plateaus was 3.44- and 3.52-fold longer compared to the WT and the Het groups (Fig. 5B–C). The probability of observing a spontaneous plateau was 10.1-fold greater in the Homozygous group relative to the WT and 8.8-fold greater than the Het group (Fig. 5D). Interestingly, there were no differences found when comparing input resistance (R_N_), action potential half-width, or AP firing rates in either the standing or running phases of the behavioral task (Supp. Fig. 3). Because the recording conditions in the in-vivo experiments were suboptimal in terms of access resistance, we repeated a set of experiments in the slice to more accurately compare the electrical properties of CA1 neurons between the WT and Homozygous groups. A comparison of single AP’s resulting from brief depolarizing current injections at the soma revealed a more hyperpolarized AP threshold (WT, –54.3 ± 0.352, Homo, –51.7 ± 0.389 mV, Fig. 6 A–B) and shorter AP duration in the WT group (WT, 1.02 ± 0.009, Homo, 1.07 ± 0.012 ms, Fig. 6C) despite similar values for resting potential, AP amplitude, maximum dV/dt, R_N_, and sag ratio (Supp. Fig. 3 A–E). In response to longer depolarizing current steps, the AP output as a function of current amplitude were similar for all but the 50 pA step, likely reflecting the difference in AP threshold (Fig. 6D–G, Supp. Fig. 3F).

**Figure 5:**
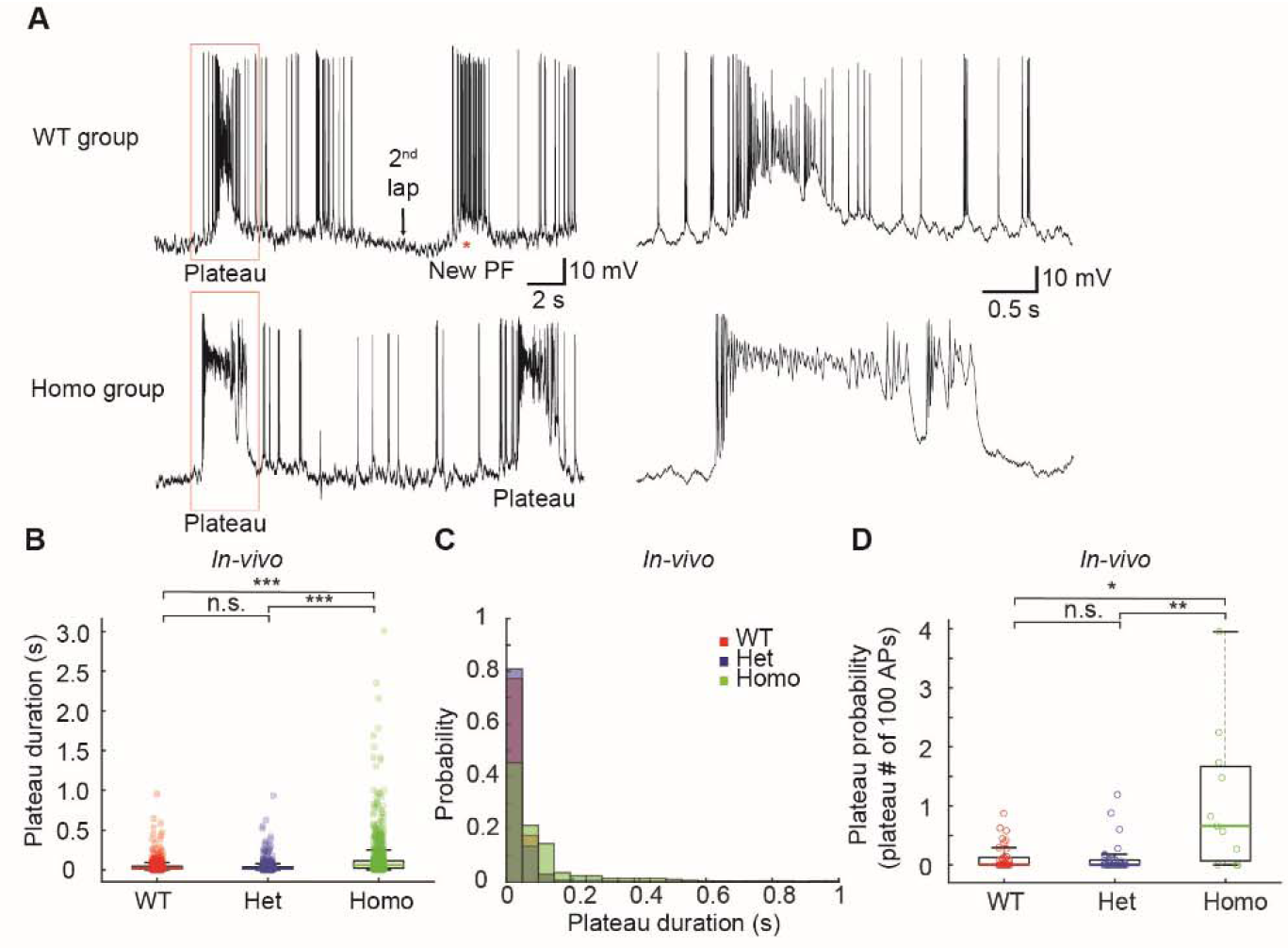
Cellular excitability comparison between WT and CaMKII T286A mutant neurons from *in-vivo* recordings. **(A)**, example traces of WT group (top) and Homo group (bottom). (**B)**, quantification of the plateau potential duration in 3 groups. All plateau potential events with a peak voltage that is more depolarized than –35 mV are used for analysis. P-values: 0.254 (WT vs. Het), 3.33e–5 (WT vs. Homo), 3.33e–5 (Het vs. Homo); n=1650, 964, and 989 for each group, respectively. (**C)**, the distribution of the duration of all the plateau potentials. The Homo group has longer duration plateau potentials. (**D)**, quantification of the plateau potential probability. Plateau probability is quantified by the amount of plateau events divided by the action potential numbers (same as described before (*27*)). Only plateau potentials with more than 50 ms duration were quantified. P-values: 0.900 (WT vs. Het), 0.0212 (WT vs. Homo), 0.00586 (Het vs. Homo). Kruskal-Wallis test with Dunn’s test for multiple comparisons was employed in panel D. Permutation test with Bonferroni correction was conducted in panel B. *P< 0.05, **P< 0.01 and ***P< 0.001; NS, not significant (P≥ 0.05). Sample sizes: n=39 (WT), n=38 (Het), n=11 (Homo).

**Figure 6:**
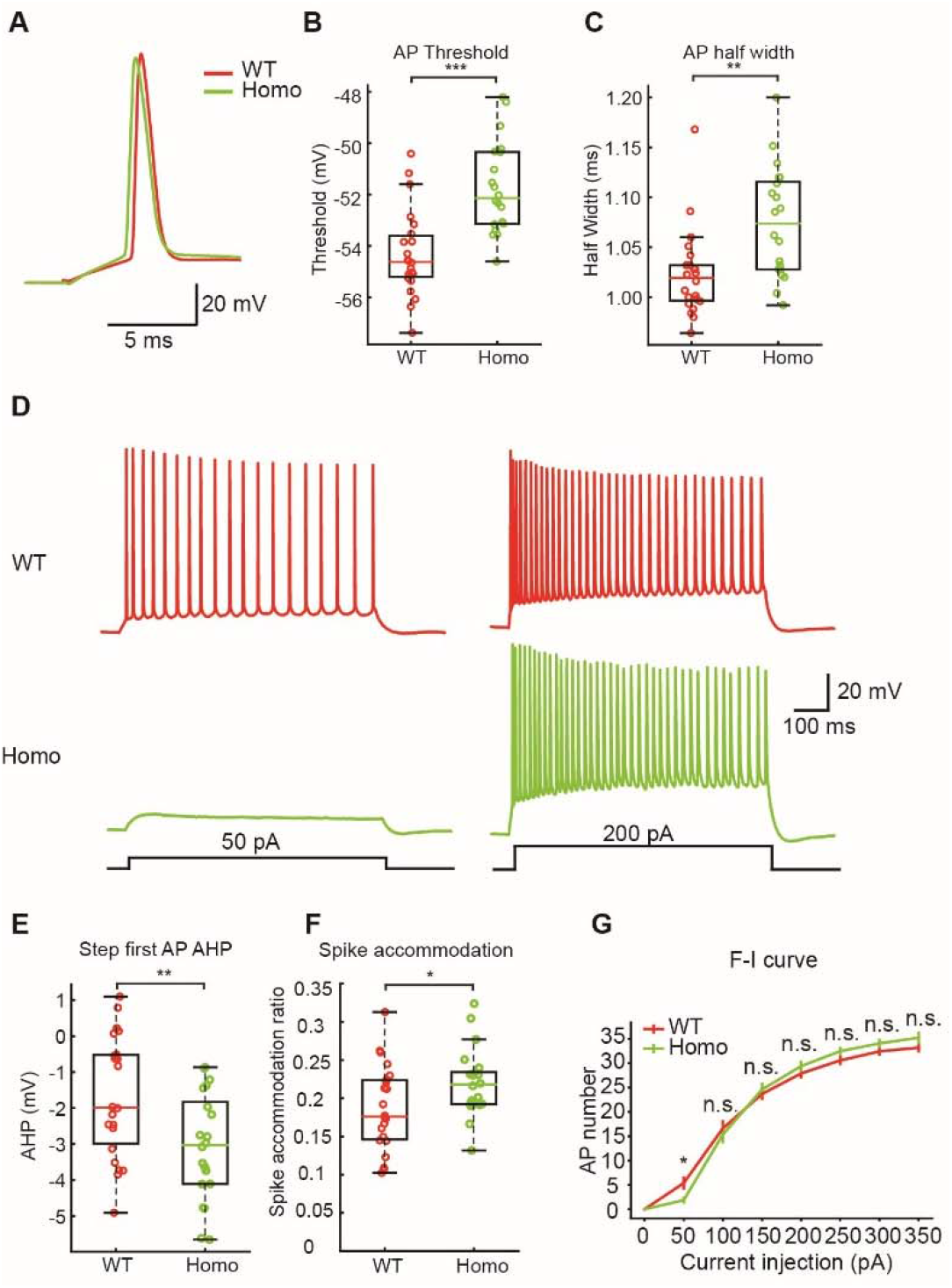
Cellular excitability comparison between WT and CaMKII T286A mutant neurons from *in-vitro* recordings. **(A)**, example trace of single action potential recorded from brain slices in WT and Homo animals. (**B)**, quantification of the action potential threshold. P-value: 2.25e–5. (**C)**, quantification of the action potential half width. P-value: 0.00170. (**D)**, example traces recorded from brain slices with step current injection in WT and Homo group mice. Only the 50 and 200 pA traces were shown here for clarity. (**E)**, quantification of the after hyperpolarization in the first action potential elicited by step current injection (0 to 350 pA in 50 pA steps). The AHP is calculated by subtracting the threshold from the minimum between the first and second action potentials. P-value: 0.00854. (**F)**, quantification of spike accommodation in WT and Homo cells with step current injection. Spike accommodation is calculated by dividing the first inter spike interval and the last inter spike interval with 350 pA current injection (see (*32*)). P-value: 0.0376. (**G)**, The comparison of the F-I curve in the WT group and Homo group. P-values: 0.0219 (rank-sum), 0.608 (t-test), 0.627 (t-test), 0.336 (t-test), 0.156 (t-test), 0.215 (t-test), 0.0607 (rank-sum). To analyze the F-I curve, every individual cell’s F-I curve was fitted to a sigmoid function. The midpoint of every fitting result was used for comparison between groups (supplementary figure 3F). Two-sample Student’s t-test is conducted for all parametric analyses. Mann-Whitney U test is conducted for all non-parametric analyses. *P< 0.05, **P< 0.01 and ***P< 0.001; NS, not significant (P≥ 0.05). Sample sizes: n=22 (WT), n=20 (Homo).

## Discussion

We have demonstrated a critical role for αCaMKII in hippocampal CA1 BTSP using a combination of in-vivo and in-vitro experimental platforms. The T286A point mutation markedly reduced the formation of PCs by synaptic plasticity induced by both spontaneous and induced plateau potentials as animals homozygous for this mutation performed a spatial navigation task. In contrast, the same interaction of plateau potentials and active synapses reliably produced PC activity in WT or Het animals (Fig. 2, 3). These results suggest that αCaMKII does not function as either an IS produced by plateau potential generation or an ET resulting from appropriate synaptic activation. Indeed, that the T286A point mutation greatly inhibited the magnitude of the plasticity instead of simply altering the shape of the resulting subthreshold V_m_ ramp (Fig. 1) suggests a role for αCaMKII signaling in the expression of BTSP.

The observation of an increased probability of spontaneous plateau potential occurrence in the homozygous T286A mice suggested the possibility of alterations in the hippocampal circuit and/or cellular excitability. If the underlying mechanisms contributing to this observation created a situation where inputs to CA1 were not reliably tuned to incoming spatial information, it could be possible that in-vivo BTSP induction would act on a set of inconsistently active synapses making any weight modifications difficult to observe. Experiments in the slice controlled for the potential lack of reliable input specificity and allowed appropriate synaptic input-plateau intervals to maximize the temporal overlap between the putative IS and ET. BTSP induction in the slice at the interval most likely to produce maximal potentiation still failed to produce significant increases in synaptic strength in slices from the T286A group but produced a 3-fold increase in EPSP amplitude in slices from WT controls. The combination of in-vivo and in-vitro experimental results suggest a role for αCaMKII outside of the IS or ET mediation, such as synaptic plasticity expression mechanisms.

The spontaneous plateau potentials observed in the T286A animals were longer in duration and occurred more frequently (Fig. 5). A comparison of cellular excitability, measured at the soma, suggested that CA1 pyramidal neurons in the T286A mice had downregulated excitatory conductances such that AP threshold was more depolarized perhaps in an attempt to compensate for the longer and more frequent plateau potentials. It is known that the probability of plateau potential generation is modulated by EC input such that silencing direct EC input to CA1 decreases plateau probability(*26*). If the lack of stable PC formation in CA1 associated with the T286A mutation somehow signals the EC to upregulate activity, this might contribute to the increase in spontaneous plateaus. In addition, the T286A mutation is believed to downregulate the activity-dependent activation of dendritic K^+^ currents, specifically those mediating the sAHP in response to suprathreshold synaptic stimulation(*29*), and inhibiting the autophosphorylation of αCaMKII blocks the homeostatic upregulation of HCN channels(*30*). Future investigations will help determine if the frequency and duration of the spontaneous plateau result from alterations to the network, uncontrolled activity-dependent changes in cellular excitability or some combination thereof.

At present, the molecular basis of ET and IS remains elusive. To address this issue, the development of new tools is crucial, particularly those capable of rapidly blocking specific molecules within seconds. Moreover, devising a method for high-throughput screening of BTSP alterations following molecular perturbations would be highly beneficial. Such advancements in technology will not only enhance our comprehension of the BTSP phenomenon but also enrich our knowledge of learning and memory in general. In summary, we have demonstrated an essential role for αCaMKII signaling in BTSP. These results should provide a starting point for future studies to uncover the molecular mechanisms involved in a form of synaptic plasticity known to underlie the formation of PCs and ultimately the cellular basis for memory formation in the hippocampus.

## Materials and Methods

### Experimental design

We hypothesize αCaMKII activity might play a role in BTSP. In order to test this hypothesis, we utilized αCaMKII T286A mutant mice for BTSP induction experiments. By examining the BTSP induction results, we can determine if αCaMKII is implicated in BTSP and whether it underlies the eligibility trace (ET) or the instructive signal (IS) or the expression mechanisms.

### Animals and procedures

All experimental procedures were approved by the Baylor College of Medicine Institutional Animal Care and Use Committee (Protocol AN-7734). All *in-vivo* experiments were performed as previously described(*5*). In-vivo whole cell recording experiments were performed on 2- to 5-month-old mice of either sex. The αCaMKII T286A mutant mouse line was provided through the courtesy of Dr. Ryohei Yasuda (Mouse Genome Informatics (MGI) id: 2158733 (*19*)).

Under deep isoflurane anesthesia, custom titanium head bars with an opening above the dorsal hippocampus were affixed to the skull using cyanoacrylate glue and dental cement. Stereotactic coordinates were employed to mark the location of the future craniotomy on the skull beneath the head bar, specifically, 1.9 mm posterior from Bregma and 1.6 mm lateral from the midline for whole-cell recordings. The recovery period post-surgery was a minimum of 7 days. After recovery, animal water intake was restricted to 1.5 ml/day. After 2 days of water limitation, animals underwent 5 days of handling and were acclimated to perform head-fixed on a linear treadmill. Animals were trained to run for a 10% sucrose water reward (∼3.5 μL per lap). The training duration for the initial three days was 20, 40, and 60 minutes, respectively. Subsequently, all animals were trained for 1-12 days until they reliably completed a minimum of 80 laps per session (60 min). Post-training, mice were provided with supplemental water to ensure a daily total of 1.5 mL. Following training, a craniotomy of approximately 0.5 mm in diameter was made above the dorsal hippocampus while the animals were under isoflurane anesthesia. The craniotomy was covered with silicone elastomer (Kwik - cast, World Precision Instruments), and animals were given approximately 24 hours to recover before recording sessions.

### Linear Treadmill

The linear treadmill consisted of a velvet fabric belt with three distinct regions of visual and tactile cues, each region occupying one-third of the belt as previously described (*5*). A custom-made lick port controlled by a solenoid valve (Lee Valve, LHQA1231220H_B) delivered the sucrose water reward. An optical sensor (Panasonic, FX-301H) detected licking. The belt’s location was reset each lap by two photoelectric sensors positioned at the beginning of the linear track. Speed was measured using a rotary encoder affixed to a treadmill wheel axle, and distance was calculated by integrating velocity within a lap. The valve, sensors, and encoder were controlled using a Bpod Finite State Machine (r1.0, Sanworks). A separate custom Arduino-based (Teensy 3.5) module was employed to control position-dependent intracellular current injection. All behavioral data were digitized at 20 kHz by a PCIe-6343, X-series DAQ system (National Instruments) using WaveSurfer software (wavesurfer.janelia.org).

### In vivo electrophysiology

Prior to conducting whole-cell patch recordings, the depth of the CA1 pyramidal layer was ascertained using an extracellular recording electrode. This procedure utilized a 1.5–3 MΩ glass pipette filled with 0.9% NaCl solution. The electrode was positioned vertically on a micromanipulator (Luigs and Neumann), and the extracellular signal was assessed using an audio monitor (A-M systems). The CA1 pyramidal layer was identified by the manifestation of theta-modulated spikes and an increased ripple amplitude. Typically, the depth of the CA1 pyramidal layer ranged from 1000 to 1300 μm below the brain surface. For whole-cell recordings, elongated-taper whole-cell patch electrodes (8–12 MΩ) were filled with a solution containing 134 mM potassium gluconate, 6 KCl, 10 HEPES, 4 NaCl, 0.3 Mg-GTP, 4 Mg-ATP, and 14 Tris-phosphocreatine. In some recordings, 0.2% biocytin was added to the intracellular solution. As the patch electrode advanced through the cortex, ∼10 psi positive pressure was applied to avert blockage. At an approximate depth of 100 μm above the pyramidal layer, positive pressure was reduced to ∼0.25 psi. Upon contact between the electrode tip and a cell, the resistance would increase reproducibly. All neural recordings were performed using a Dagan BVC-700A amplifier in current clamp mode and were digitized at 20 kHz by a PCIe-6343, X series DAQ system (National Instruments) employing WaveSurfer software (wavesurfer.janelia.org). Bridge balance was adjusted to compensate for series resistance. Recordings with a series resistance exceeding 60 MΩ were excluded from subsequent analyses.

### In vitro electrophysiology

Horizontal hippocampal slices of 400 μm thickness were obtained from 6- to 12-week-old male and female mice using a Leica Vibratome VT1200S. Animals were anesthetized with isoflurane and ketamine/xylazine injection, followed by intracardial perfusion with an ice-cold cutting solution containing (in mM) 205 sucrose, 25 NaHCO_3_, 2.5 KCl, 1.25 NaH_2_PO_4_, 1 CaCl_2_, 7 MgCl_2_, and 10 glucose. The slices were incubated in standard aCSF (below) for 40 minutes at 35°C before being maintained at room temperature. Whole-cell current-clamp recordings were conducted at 33-34°C in aCSF perfused into a submerged recording chamber at a rate of 2 ml/min. The aCSF composition varied depending on the experiment and contained (in mM): 125 NaCl, 25 NaHCO_3_, 2.5 KCl, 1.25 NaH_2_PO_4_, 2 CaCl_2_, 1 MgCl_2_, 16 glucose (for BTSP experiments); 125 NaCl, 25 NaHCO_3_, 3 KCl, 1.25 NaH_2_PO_4_, 1.3 CaCl_2_, 1 MgCl_2_, 16 glucose (for all other experiments). All aCSF solutions contained fresh 3 mM sodium pyruvate and 1 mM ascorbic acid and were constantly bubbled with 95% O_2_ and 5% CO_2_.

Cells were visualized using a Zeiss Examiner Z1 microscope with a water-immersion lens (63x, 1.0 NA, ZEISS) using Dodt contrast. Whole-cell patch recordings were performed using a Dagan BVC-700 in current clamp mode, analog-filtered at 1 kHz and digitized at 50 kHz using a PCIe-6343, X series DAQ system (National Instruments), controlled by Neuromatic software (*31*). Patch electrodes (5–7 MΩ) were filled with filtered internal solutions, varying based on the experiment. For BTSP experiments, the solution contained (in mM): 130 Cs-methanesulfonate, 6 KCl, 10 HEPES, 4 NaCl, 4 Mg-ATP, 0.3 Tris-GTP, and 14 Tris-phosphocreatine. For all other experiments, the solution contained (in mM): 134 potassium gluconate, 6 KCl, 10 HEPES, 4 NaCl, 0.3 Mg-GTP, 4 Mg-ATP, 14 Tris-phosphocreatine, and 0.2% biocytin.

Series resistance was maintained between 18–45 MΩ when using Cs^+^ to optimize cell health as previously described (*5*). EPSPs were induced by extracellular stimulation (0.1 ms, 0.1–0.5 mA) of axons in the *stratum radiatum* region using a platinum-iridium microelectrode (0.5 MΩ, WPI) positioned 100–300 μm from the recorded cell. Input resistance was monitored using a 150 ms, – 30 pA current injection following the EPSP test stimulation. For BTSP experiments, gabazine (SR 95531, 2 μM) and CGP 55845 (50 nM) were added to the aCSF solution, and the CA3 region was removed to prevent epileptiform activity.

### Cell inclusion criteria for in-vivo recordings

A subset of cells was included for analysis (Supplementary Figure 4), based on the following criteria: (1) The cell’s afterhyperpolarization (AHP) was quantified by determining the mode of all action potential AHPs, with the requirement that it should exceed –5 mV. (2) The mean squared error of the membrane potential (Vm) distribution fit to a Gaussian function should be more than 4.5e–6. Criteria 1 and 2 were designed to identify potential interneurons (*28*). (3) The animal’s locomotion during the induction laps should not include more than 10 stops (excluding stops at the reward site to consume the reward). (4) The licking probability for areas outside the reward zone should be less than 0.1. The reward zone was defined as 12.6 cm before the reward and 14.4 cm after the reward. Criteria 3 and 4 aimed to exclude cells correlated with poor animal behavior. (5) The action potential (AP) amplitude should be greater than 35 mV relative to threshold. The AP amplitude is indicative of recording quality, with higher AP amplitudes signifying better recording quality. This criterion was established to exclude cells with poor recording quality. (6) The most hyperpolarized part of the ramp (5-bin average) should not exceed 8 mV. This criterion was set to exclude cells in which the BTSP induction was too close to the initiation of the recording when cells are transiently hyperpolarized. (7) Cells from the CA3 region were also excluded. As a result of the selection process, 88 out of 125 recorded cells (70.4%) were included. However, quantification of the entire 125 cells revealed similar results (both the behavior and the BTSP) to the selected 88 cells (Supplementary Figure 4, 5A–5D), suggesting that the cell selection process did not affect the final conclusions.

### The correlation of behavior and BTSP

Animal behavior quantified as licking and running has no correlation (or very weak correlation) to the BTSP ramp formation (Supplementary Figure 5E, F), suggesting the difference in the behavior of CaMKII homozygous mutant group does not affect the BTSP induction experiment.

### Data analysis

For the analysis of the ramp of depolarization, action potentials were eliminated by removing all data points 1.5 ms before and 3.5 ms after a designated threshold value (the maximum second derivative with respect to voltage) and low-pass filtered (<3Hz) as described previously (*5*). The subthreshold membrane potential was spatially binned into 100 bins, each approximately 1.8 cm in size. The average V_m_ from five laps preceding BTSP induction and five laps following the induction was calculated to determine the before trace and after trace. The ramp was computed by subtracting the before trace from the after trace, subsequently converting it to the time domain based on the fastest running speed during the five induction laps. To quantify the peak of the ramp, the membrane potential from 1 s before running and 1 s after the animal completed running was incorporated into the ramp (with 20 time bins added before the 100 spatial bins and 20 time bins after the spatial bins). The average V_m_ recorded in the 20 time bins preceding the animal’s initiation of movement served as the baseline for the ramp and was subtracted from all values to set the baseline V_m_ to 0 mV. The ramp peak was defined as the maximum value of the baseline-subtracted subthreshold V_m_, and the ramp’s area was calculated from the integral of the subthreshold V_m_.

Spike frequency accommodation was analyzed by examining action potential (AP) trains resulting from a 350 pA current injection, as the number of action potentials in the WT and Homo groups were not statistically different at this current injection intensity (Figure 6F). Spike accommodation was quantified by dividing the first interspike interval by the last interspike interval.

Animal behavior was analyzed by quantifying the animal’s licking behavior and running speed. The animal’s running speed was calculated using the average of all the recorded laps of the spatially binned running speed. Grouped data shown in Figure 2E are the mean and SEM of each recorded cells in 3 groups. The spatial bin that has the minimum average running speed was used for comparison between the 3 groups as shown in Figure 2F. The licking probability was calculated for each spatial bin, with a probability of 1 if a lick was detected and 0 if no lick was detected. The relative licking probability increase in the reward zone was calculated as the averaged licking probability within the reward zone minus the averaged licking outside the reward zone, divided by the averaged licking outside the reward zone, as defined by the following equation:

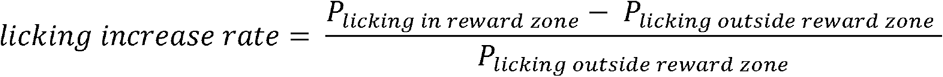

The reward zone was defined as 12.6 cm before and 14.4 cm after the reward. This method was employed for quantifying licking behavior. Ideal licking behavior should exhibit high licking probability within the reward zone and very low licking probability outside of the reward zone.

### Statistical methods

Sample sizes were not predetermined using statistical methods. Data analysis was performed using two-sample Student’s t-test or one-way ANOVA with Bonferroni correction, as indicated in the figure legends. Non-parametric tests employed Kruskal-Wallis test with Dunn’s test for multiple comparisons. Normality of the data was assessed using the Shapiro-Wilk parametric hypothesis test, using a MATLAB script (Ahmed BenSaïda, (https://www.mathworks.com/matlabcentral/fileexchange/13964-shapiro-wilk-and-shapiro-francia-normality-tests)), and by visually examining the quantile-quantile plot of the data. For datasets with large sample sizes (Figure 5B), a permutation test with Bonferroni correction was conducted to avoid type I errors. The permutation test was performed using a MATLAB script (Laurens R Krol, (https://github.com/lrkrol/permutationTest)).

## Acknowledgments

We would like to thank members of the Magee lab for helpful discussions and technical assistance. We would also like to thank Evan Campbell and Brennan Sullivan for their comments on the manuscript. We also thank Dr. Ryohei Yasuda for providing the CaMKII T286A mutant mouse line.

## Author contributions

Conceptualization: JCM, KX

Methodology: JCM, RAC, KX,

YL Investigation: KX

Visualization: KX

Supervision: JCM, RAC

Writing—original draft: KX, RAC

Writing—review & editing: KX, YL, RAC, JCM

## Data and materials availability

All data, code, and materials used in the analyses are available upon request.

