## Supplemental figure 1-5 for "A critical role for CaMKII in behavioral timescale synaptic plasticity in hippocampal CA1 pyramidal neurons"

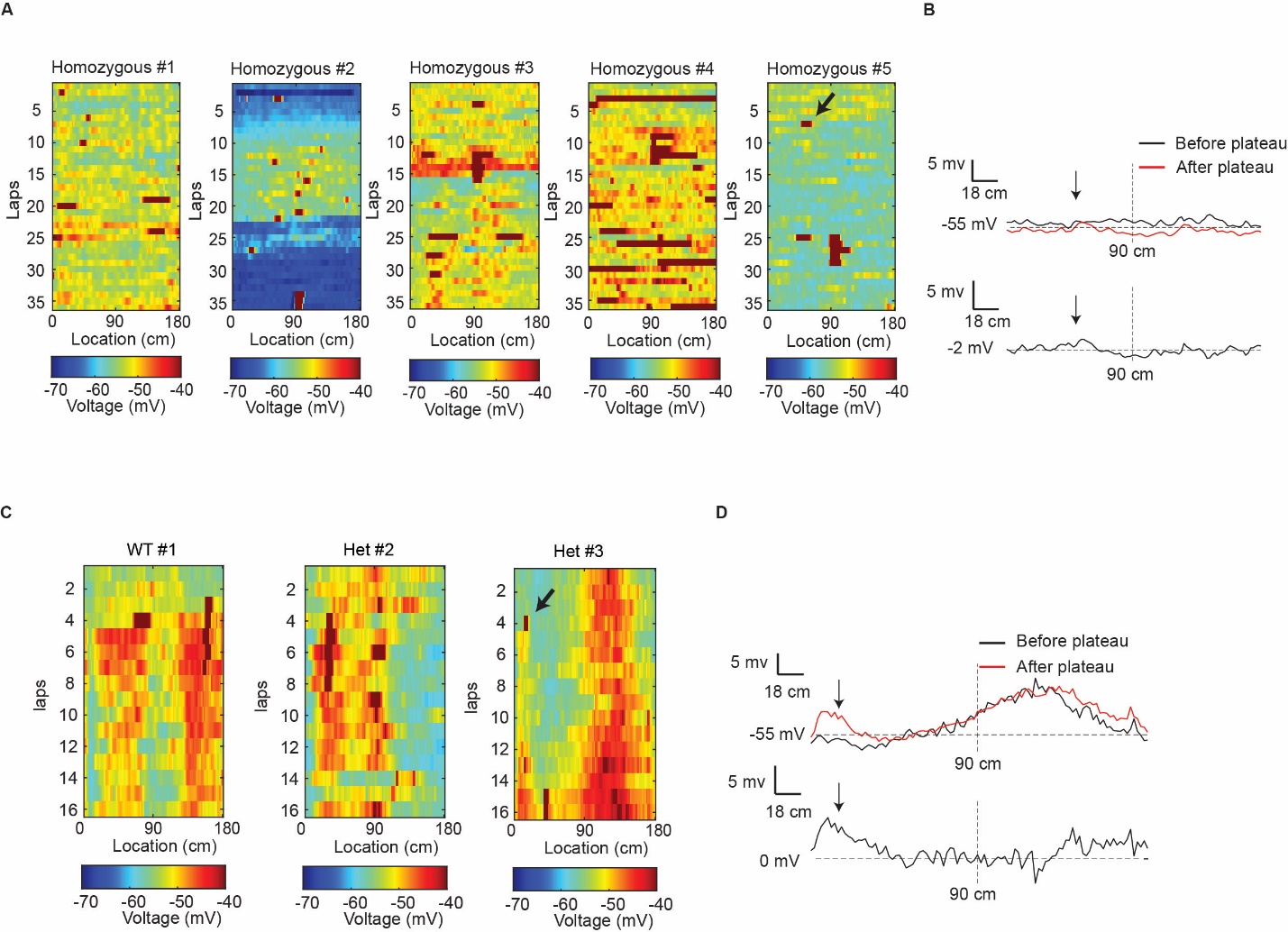

Fig. S1.

**Spontaneous plateau potential does not induce place field formation in the CaMKII homozygous mutant mice**

**(A)**, 5 example cells show the spontaneous plateau potentials in the CaMKII homozygous mutant mice do not induce place field formation. Note that the black arrow indicates a spontaneous plateau event. (**B)**, Plot of the subthreshold membrane potential before and after the spontaneous plateau event. This Plot corresponds to the plateau event marked by the black arrow in panel A. Plateau location is marked on top of the traces by arrowheads. (**C)**, 3 examples of spontaneous plateaus in both the WT and the Het group. (**D)**, Plot of the subthreshold membrane potential before and after the spontaneous plateau event. This Plot corresponds to the plateau event marked by the black arrow in panel C. Plateau location is marked on top of the traces by arrowheads.

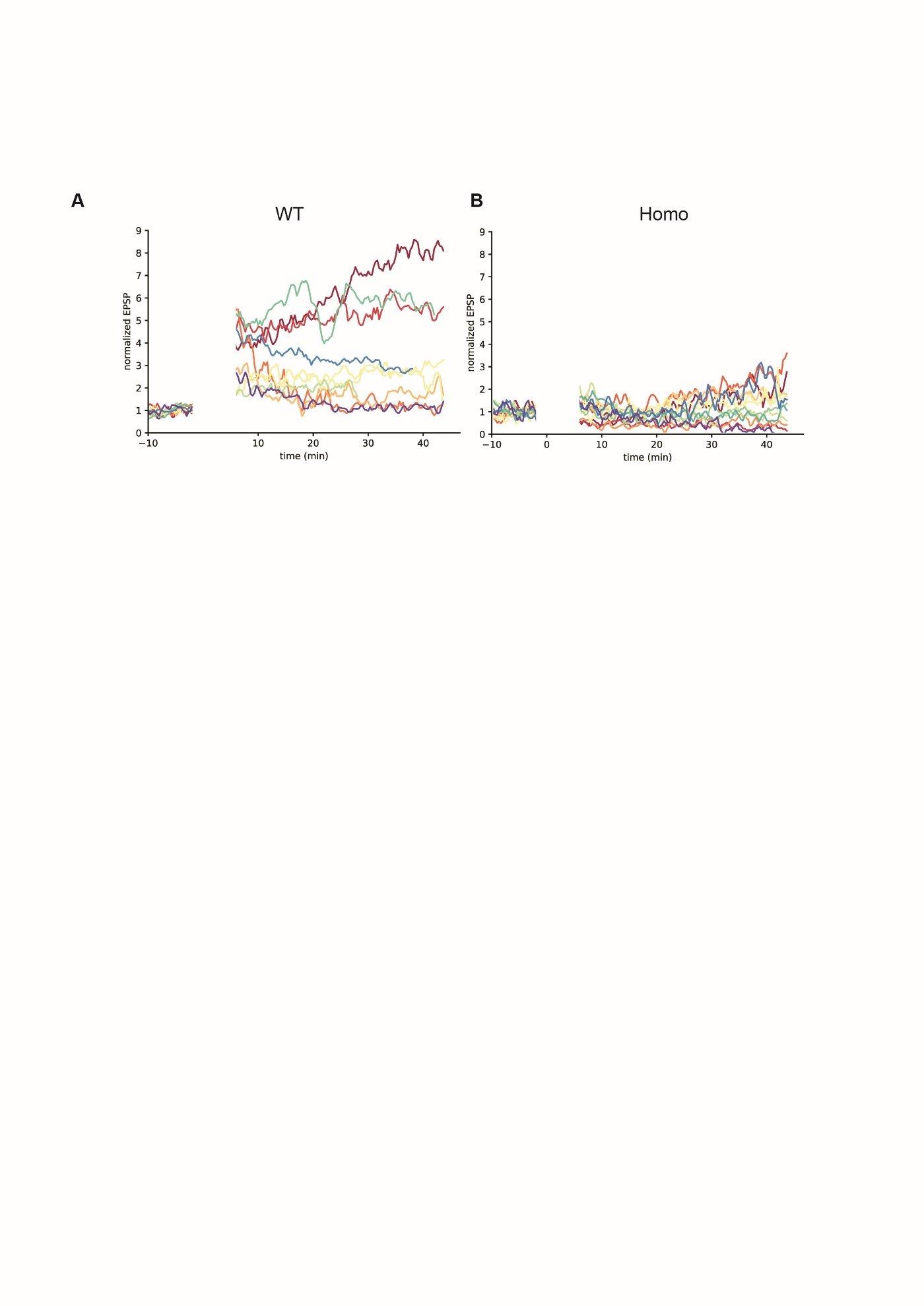

Fig. S2.

**All neurons tested for BTSP in slice**

**(A)**, BTSP induction causes EPSP amplitude to increase in cells from WT group. (**B)**, BTSP induction is less effective in the cells from Homo group. n=10 for WT, n=12 for Homo.

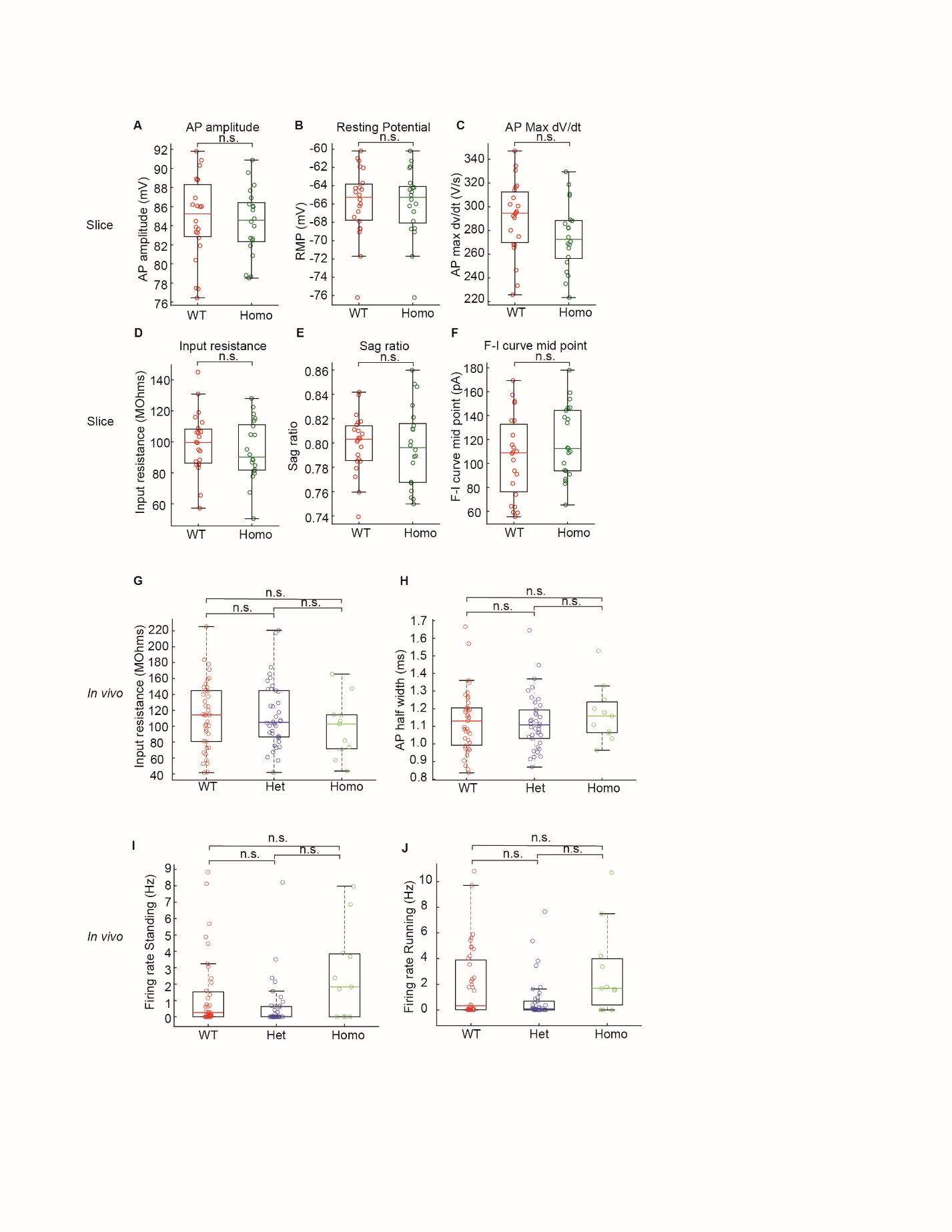

Fig. S3.

**A comparison of membrane properties between the CaMKII T286A and WT neurons**

**(A)**, quantification of the action potential amplitude in WT and Homo groups cells recorded from brain slices. P-value: 0.713. (**B)**, quantification of the resting potential in WT and Homo groups cells recorded from brain slices. P-value: 0.895. (**C)**, quantification of the max dv/dt in action potential in WT and Homo groups cells recorded from brain slices. P-value: 0.0912. (**D)**, quantification of the input resistance in WT and Homo groups cells recorded from brain slices. P-value: 0.458. (**E)**, quantification of the sag ratio in WT and Homo groups cells recorded from brain slices. P-value: 0.840. (**F)**, quantification of the mid-point in the F-I curve fitted to a sigmoid function. P-value: 0.196. (**G)**, comparison of input resistance recorded in-vivo in WT, Het and Homo groups. P-value: 1.00 for WT and Het; 0.911 for WT and Homo; 0.402 for Het and Homo. (**H)**, comparison of action potential half width recorded in-vivo in WT, Het and Homo groups. P-value: 0.995 for WT and Het; 0.845 for WT and Homo; 0.765 for Het and Homo. (**I)**, quantification of the standing firing rate of 5 laps before BTSP induction. P-value: 0.106 for WT and Het; 0.903 for WT and Homo; 0.130 for Het and Homo. (**J)**, quantification of the running firing rate of 5 laps before BTSP induction. P-value: 0.294 for WT and Het; 0.679 for WT and Homo; 0.112 for Het and Homo. Two sample student’s t-test was conducted in all slice experiment analysis. One-way ANOVA with Bonferroni correction was conducted in all parametric *in-vivo* experiment statistical analysis. Kruskal Wallis test with Dunn’s test for multiple comparisons in non-parametric analysis. *P< 0.05, **P< 0.01 and ***P< 0.001; NS, not significant (P≥ 0.05). n=22 for WT, n=20 for Homo in slice experiment. n=39 for WT, n=38 for Het, n=11 for Homo in in-vivo experiments.

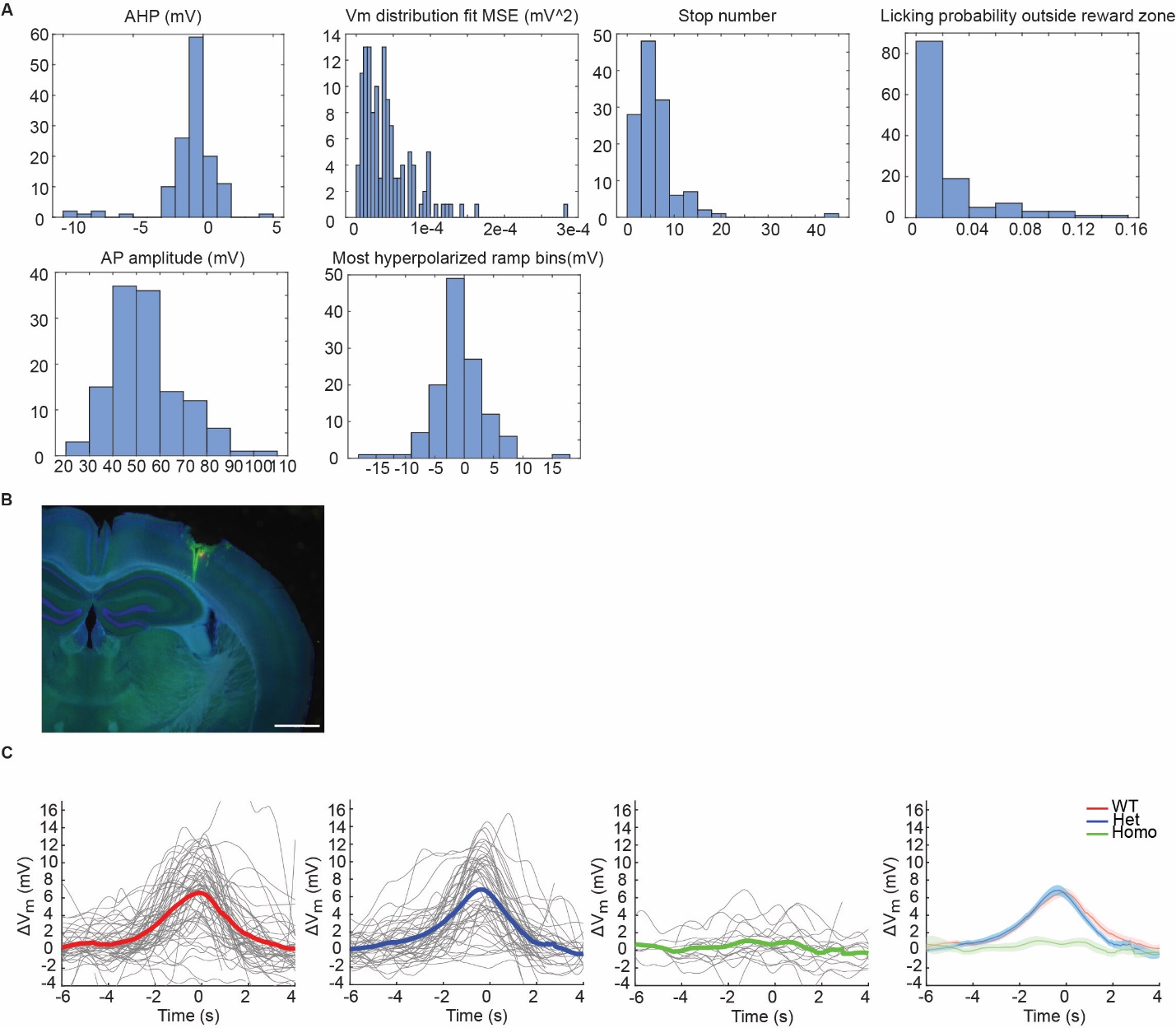

Fig. S4.

**Cell selection does not affect BTSP ramp analysis results**

**(A)**, The distribution of 6 parameters that are used in cell selection criteria (see Methods). Cell selection criteria contains 7 standards: 1. The cell’s AHP (mode of all action potential’s AHP) should be > –5 mV. 2. The mean squared error of V_m_ distribution fit to gaussian function should be more than 4.5e–6. The purpose for standards 1 and 2 is to exclude potential interneurons (see(*1*)). 3. The animal’s average stop number during the induction period should be less than 10 times (excluding the stop at the reward site to consume the reward). 4. The licking probability for outside reward zone should be less than 0.1. reward zone is defined as 12.6 cm before reward and 14.4 cm after reward. 5. The AP amplitude should be > 35 mV. In *in-vivo* recordings, the AP amplitude is a good indication of the recording quality, with higher AP amplitude, the recording quality is better. Here the standard is to exclude cells that has poor recording quality. 6. The most hyperpolarized part of ramp (5 bins average) should not be > 8 mV. This standard is to exclude cells that have BTSP induction too close to the beginning of the recording. 7. Cells from CA3, see panel B. (**B)**, example of cell from CA3 region. scale bar, 1 mm. **C**, plot of ramps from all cells collected in WT Het and Homo groups. Note that the shape of the ramp is very similar to the cells selected for analysis.

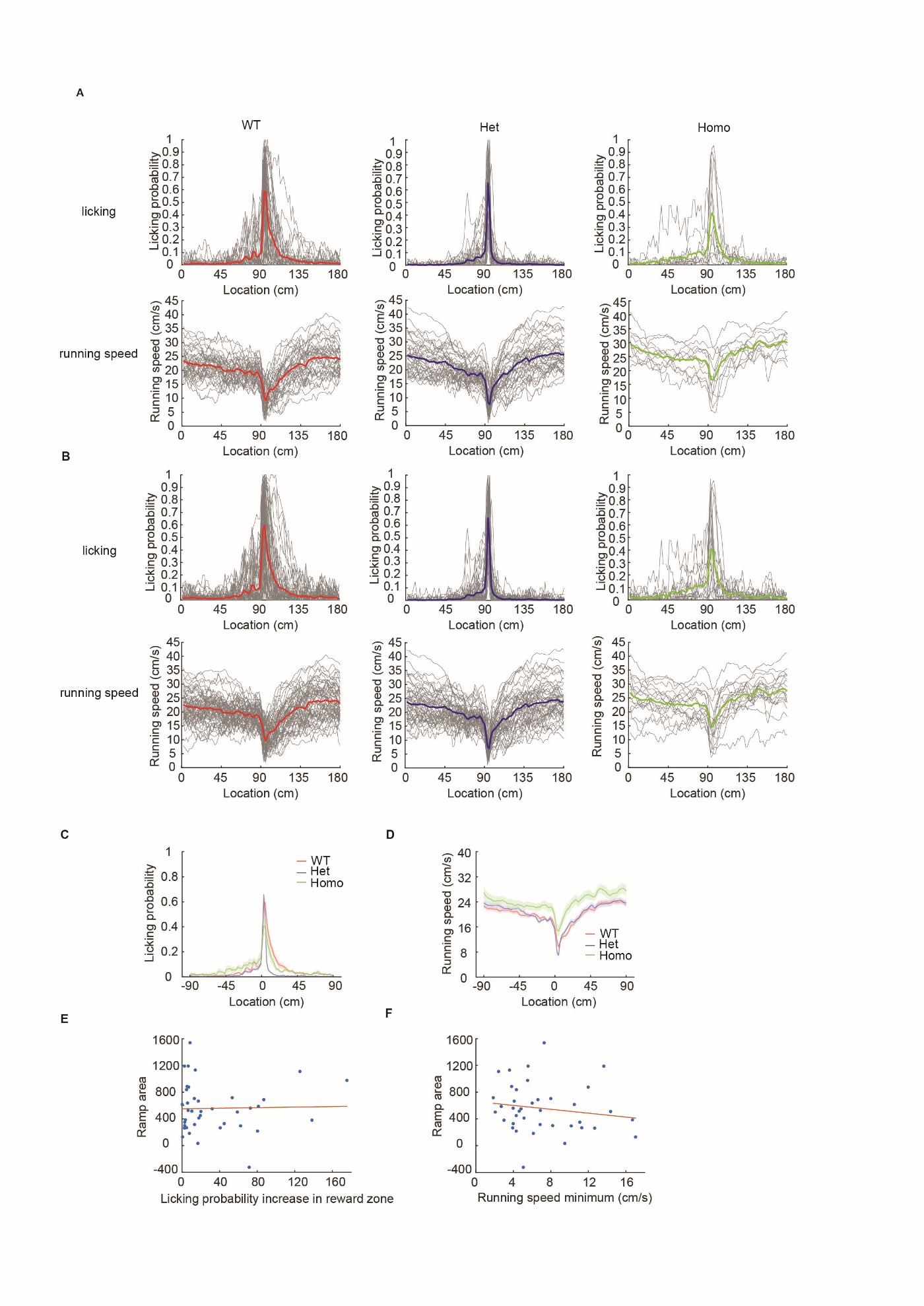

Fig. S5.

**Cell selection does not affect the behavior analysis results**

**(A)**, The licking and running traces from the subset of cells that have been selected using the inclusion criteria (N=88, see methods). (**B)**, The licking and running traces from all cells (N=125) that have been recorded. (**C)**, The averaged traces of the licking probabilities in WT, Het and Homo groups from all cells recorded. The central line and shaded background represents mean ± SEM, respectively. (**D)**, The averaged traces of the running speeds in WT, Het and Homo groups from all cells recorded. (**E)**, The correlation between the animal behavior (licking) and the BTSP ramp. The regression showed that the Licking was not (or very weakly) correlated with the BTSP ramp. R-square for regression: 5.67e–4. (**F)**, The correlation between the animal behavior (running) and the BTSP ramp. The plot showed the BTSP ramp area plotted against the minimum running speed. R-square for regression: 0.0260. n=58 for WT, n=50 for Het, n=17 for Homo in all cells. N=39 for WT, n=38 for Het, n=11 for Homo in selected cells.
